# The Parkinson’s disease-associated A53E mutation decreases alpha-synuclein aggregation and phosphorylation *in vivo*

**DOI:** 10.64898/2026.10.29.762882

**Authors:** WL Irving, S Meredith-Andrews, KT Nguyen, KL Schainker, NL Brockway, VK Unni, TA. Weissman

## Abstract

Abnormal α-synuclein aggregation is the defining feature of Parkinson’s disease (PD) and related synucleinopathies. Rare disease-causing α-synuclein mutations either accelerate or slow fibrillization *in vitro*, however their effects *in vivo* are less well understood. Interestingly, the A53E mutation slows fibrillization *in vitro* but causes a more virulent form of PD. We use an *in vivo* model to examine the aggregation dynamics of disease-causing A53E and E46K α- synuclein mutations. Fluorescence recovery after photobleaching within presynaptic terminals shows that A53E-α-synuclein is significantly more mobile than E46K- or wild-type (WT)-α- synuclein. Longitudinal analysis *in vivo* reveals that E46K- and WT-α-synuclein become progressively less mobile over days, while A53E maintains high mobility, suggesting delayed progression toward aggregated states. We also find that A53E-α-synuclein lacks detectable disease-relevant phosphorylation at serine-129, unlike E46K-and WT-α-synuclein. To mechanistically test whether this unique lack of phosphorylation drives its high mobility, we genetically modified A53E-α-synuclein to mimic S129 phosphorylation, and show that this manipulation indeed decreases its mobility, directly implicating the phosphorylation status of this residue in mobility.

## Introduction

Parkinson’s disease (PD) is a chronic, progressive neurodegenerative disorder defined by the accumulation of Lewy bodies, which are composed largely of aggregated α-synuclein protein found in the cytoplasm of neurons^1^. In healthy neurons, the 140-amino-acid α-synuclein protein is largely soluble and localized primarily in presynaptic terminals, where it regulates neurotransmitter release and vesicle cycling^2,3^. Beyond the synapse, α-synuclein also plays an essential role in the nucleus, where its absence increases double-strand DNA breaks and reduces DNA repair capacity^4^. The formation of Lewy bodies in disease may thus lead to a toxic gain-of-function of the aggregated protein in the presynaptic terminal or other cellular locations^5^, a potential loss-of-function via nuclear mechanism^4,6,7^, or some combination of these processes. However, despite decades of research, the biological mechanisms underlying α-synuclein aggregation and Lewy body formation remain poorly understood. A better understanding of these processes is critically important, since there are currently no treatments to halt the progression of PD or other Lewy body disorders.

Familial forms of PD in patients who carry autosomal dominant mutations offer one clear avenue for investigating the mechanisms underlying α-synuclein toxicity. In these cases, a single amino acid change significantly impacts aggregation and initiates disease. There are six well characterized familial α-synuclein mutations that cause PD: A30P^8^, E46K^9^, H50Q^10^, G51D^1^^1^, A53E^1^^2^, and A53T^13^. In patients with these mutations, the process by which each single amino acid change causes neurodegeneration is not well understood. Interestingly, three of the six known mutations, E46K, A53T, and H50Q, have been shown *in vitro* to accelerate aggregation and promote fibril formation^14–16^, while the others, A53E, A30P, and G51D, reduce or slow fibrillization^17–20^. This diversity suggests that each mutation may drive neurodegeneration through a distinct pathogenic mechanism and that the fibrillar form itself may not be causing toxicity, highlighting the need to study how these mutations affect aggregation *in vivo* within the cellular environment of neurons over time. What makes these mutations even more interesting is that they cause varying degrees of disease age onset, severity, and death. For example, families carrying the A53E mutation have an earlier age of onset and more aggressive PD progression compared to many other familial variants^21^, possibly indicating that this mutation, which surprisingly decreases aggregation, causes particularly severe pathogenic effects.

Although Lewy bodies are the pathological hallmark of PD, growing evidence suggests that small, soluble α-synuclein oligomers may be the most toxic species of aggregate^5,22^. Pathological α-synuclein aggregation is thought to progress from soluble monomers to small oligomeric species, which assemble into larger fibrils that eventually accumulate to form mature Lewy bodies^23^. *In vivo* studies have shown that neuronal dysfunction can occur in the absence of large, later-stage inclusions, further supporting the idea that early-stage oligomeric species could be the key drivers of neurotoxicity^5,24^. Mutations that favor smaller aggregates, like A53E^18,19,21^, could thus be more toxic than those that promote larger fibrils. Such a mechanism could help explain how mutations that both increase and decrease fibrillar aggregation all lead to the same disease. A detailed comparison of the disease forms of α-synuclein at the cellular level, ideally over time in an *in vivo* system, could provide insights into whether and how each mutation influences aggregation and disease.

Here we used *in vivo* imaging approaches to quantify the mobility of A53E, E46K, and wild-type (WT) forms of α-synuclein, and to investigate their propensity to aggregate over time in the zebrafish nervous system. Zebrafish offer a powerful opportunity to investigate α-synuclein-associated pathology within a transparent, intact, living vertebrate nervous system, and have been used to identify α-synuclein and other neurodegenerative mechanisms and potential therapeutic interventions *in vivo*^25–29^ (reviewed in ref ^30^). We measured mobility at multiple different synaptic locations within individual axonal arbors *in vivo* using fluorescence recovery after photobleaching (FRAP), a technique that has been established to estimate protein aggregation state^27,31–34^. Our *in vivo* platform allowed us to not only measure the mobility of various forms of α-synuclein at one individual time point, but also to measure how aggregation progressed over time in the living nervous system. Using this approach, we demonstrate that A53E-α-synuclein exists in a more mobile state than E46K- and WT-α-synuclein, and that it remains mobile over days *in vivo,* while both E46K-α-synuclein and WT-α-synuclein become progressively less mobile. We also followed the same individual neurons over four days *in vivo* and found α-synuclein protein mobility to decrease over time within the cytoplasm for each condition, except for A53E-α-synuclein, which remains relatively mobile. Corroborating these results, the level of *in vivo* phosphorylation at serine-129 (S129), a known disease-associated post-translational modification, is abolished in A53E-α-synuclein-expressing neurons at 4 dpf, in contrast to both E46K- and WT-α-synuclein-expressing neurons. To test the possibility that the increased mobility of A53E-α-synuclein is due to a lack of phosphorylation at S129, we created an additional form of α-synuclein that contains the A53E mutation along with a phosphomimetic modification at S129 (A53E/S129D). Indeed, this modification to A53E-α-synuclein partially reduced its mobility to levels closer to WT-α-synuclein. Taken together, these results show that the disease-causing A53E mutation in α-synuclein maintains the protein in a more mobile state *in vivo* for a prolonged period of time as compared to other forms of α-synuclein, and that this may occur via a concomitant decrease in S129 phosphorylation. Our *in vivo* longitudinal approach provides a powerful model for dissecting the pathological mechanisms that underlie α-synuclein aggregation.

## Results

### Alpha-synuclein disease mutations A53E and E46K can be expressed in zebrafish and localize similarly to wild-type α-synuclein

To begin to assess the effects of disease-associated mutations A53E and E46K on α-synuclein’s propensity to aggregate *in vivo,* we transiently expressed human WT, A53E, and E46K forms of α-synuclein with a C-terminal Green Fluorescent Protein (GFP) tag in zebrafish larvae (Figure 1). Zebrafish provide an accessible vertebrate model for studying protein aggregation in neurodegenerative disease by enabling direct live imaging of protein behavior in intact neurons within a visible, living nervous system^27,28,30^. To drive expression in the central and peripheral nervous systems, we used the neuroD promoter, a pan-neuronal transcription factor expressed from zebrafish development through adulthood^35^. Previous work has shown that human WT-α- synuclein expresses and functions normally in zebrafish motor neurons, localizing to presynaptic terminals^27,28,36^ and undergoing phosphorylation via endogenous zebrafish Polo-like kinases^27^. Similar to WT-α-synuclein, A53E- and E46K-α-synuclein demonstrated a characteristic punctate presynaptic terminal localization (Figure 1A-D), while free GFP was more consistently distributed throughout neurons, with more even expression throughout the axons (Figure 1E-F; also previously quantified in ref ^27^. No significant differences in axonal morphology were detected across conditions, including the size and spacing of presynaptic terminals (Figure 1G- H). These results suggest that disease forms of human α-synuclein express normally within zebrafish neurons, localize to synaptic terminals, and provide a suitable model for studying their *in vivo* dynamics.

**Figure One.**
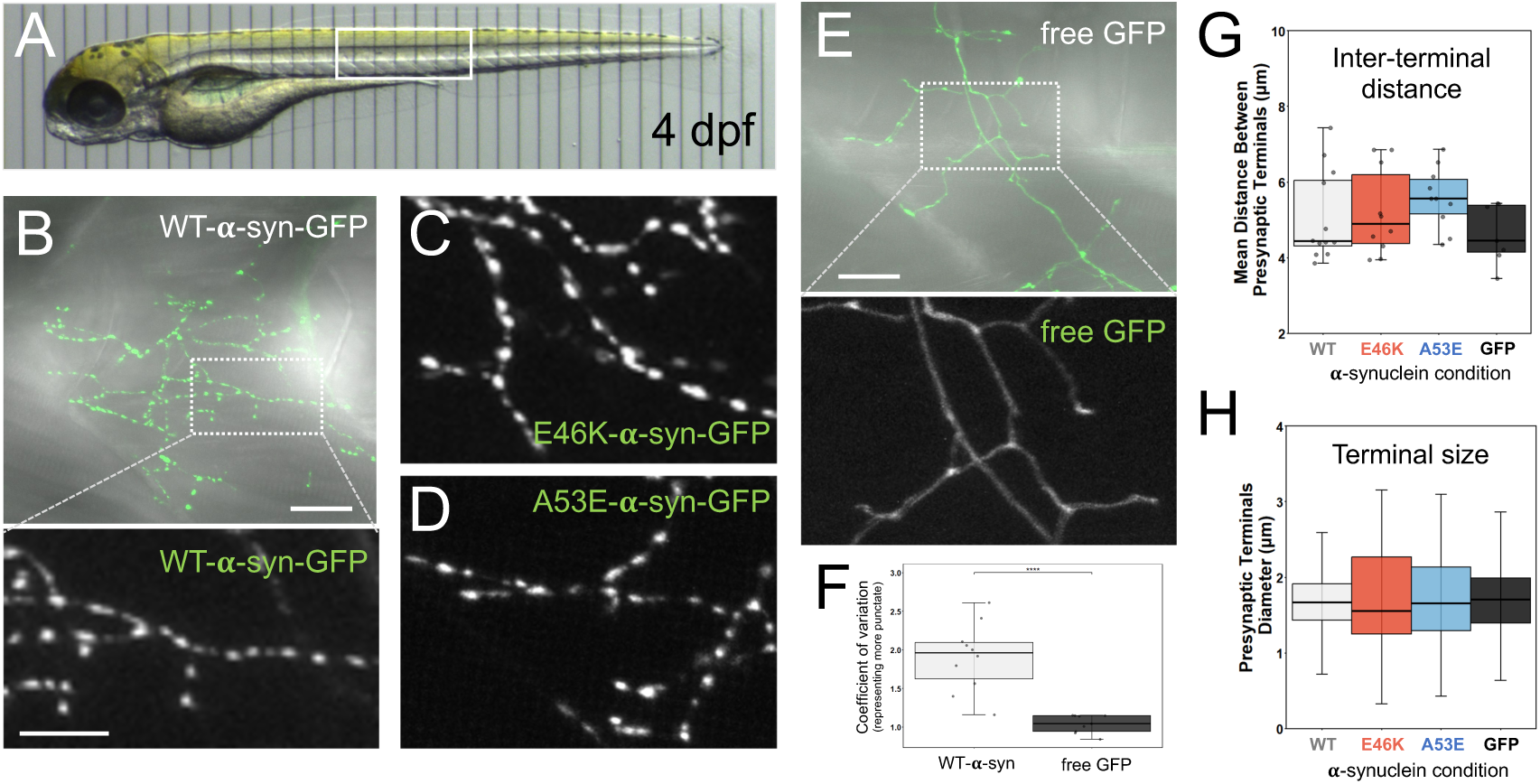
Human α-synuclein disease mutations E46K and A53E express in zebrafish. A) A representative living zebrafish larva at 4 days post fertilization (dpf) mounted on a micrometer displays the transparency of the nervous system and general region imaged (white box). Tick marks are 100 um. B) Neuron expressing WT-α-synuclein-GFP, shown overlaid onto transmitted light channel to demonstrate trunk morphology and *in vivo* context. Dashed box indicates synaptic regions shown at a higher magnification in corresponding lower panel (GFP channel). C) Neuron expressing E46K-α-synuclein-GFP. D) Neuron expressing A53E-α-synuclein-GFP. D) Neuron expressing free GFP, shown overlaid onto transmitted light channel, with dashed box indicating region magnified below (GFP channel). Images in B-E are maximum intensity projections of Z-stacks with the following depths: B) 46.2 μm; C) 40.6 μm; D) 18.9 μm; E) 63.7 μm. Scale bar is 20 μm in B and E upper panels, 10 μm in all magnified GFP panels. F) Coefficient of intensity variation calculated along axons expressing either WT-α-synuclein-GFP (1.9±0.4) or free GFP (1.0±0.1), with higher variation representing more punctate labeling, and lower variation representing more even expression along axon (Mann-Whitney test, p<0.0001). Dots represent individual neurons quantified. G) Quantification of distance between synaptic terminals for neurons expressing WT-, E46K-, or A53E-α-synuclein, or free GFP-. Mean distance between terminals was WT=5.1±1.2 μm, n=50 filaments from 12 neurons across 10 larvae; E46K=5.2±1.2 μm, n=37/10/10; A53E = 5.6±0.8 μm, n=26/10/9; free-GFP=4.63±0.79, n=19/7/5; ANOVA p=0.17. Dots represent individual neurons (each with many filaments averaged). H) Quantification of terminal size. Mean terminal size was WT=1.63±0.47 μm, n=513 terminals; E46K=1.70±0.63 μm, n=580; A53E=1.71±0.58 μm, n=317; free-GFP=1.67±0.42, n=600; ANOVA p=0.06.

### A53E- α-synuclein is more mobile than WT- and E46K-α-synuclein at 4 days

We next tested whether human disease mutations affect α-synuclein protein aggregation *in vivo* by measuring the mobility of disease-relevant forms of α-synuclein and comparing them to WT. We conducted *in vivo* FRAP experiments in the synaptic terminals of zebrafish motor neurons within living zebrafish expressing either WT-, A53E-, or E46K-α-synuclein, or free GFP (Figure 2A-C and SF1). We chose to focus these experiments at 4 days post fertilization (dpf), when motor neurons are significantly developed and display strong expression of α-synuclein-GFP. In FRAP, quantifying the fluorescence recovery over time following the bleaching of a small region of interest is used to estimate the speed of fluorescent proteins moving into the region. Small, unaggregated forms of a protein move more quickly than larger, aggregated forms^31,32,37^. Tau represents the time it takes for moving fluorescent protein to recover ∼63.3% of its final fluorescence, while the immobile fraction (IF) is calculated from the amount of fluorescence that never recovers during the course of the experiment, representing immobile, aggregated protein^33,38,39^. Both tau and IF values are predicted to increase when protein is in a more aggregated state. As expected, consistent with previous findings^27,34^, both tau and immobile fraction for WT-α-synuclein were significantly larger than free GFP at 4 dpf (Supplemental Figure 1). This is in line with evidence that expression of WT-α-synuclein leads to some aggregation, both *in vitro*^36,40^ and *in vivo*^27,28,34,39,41^, especially when it is present at high levels, for example in human duplication and triplication disorders that cause PD^42,43^. We next compared the fluorescence recovery curves, immobile fraction, and tau measurements among WT-, A53E-, and E64K-α-synuclein conditions (Figure 2D-F). Interestingly, we found that A53E-α-synuclein was significantly more mobile than both WT and E46K variants, suggesting a decreased aggregation state *in vivo*. The immobile fraction of A53E-α-synuclein was significantly lower than E46K-α-synuclein (Figure 2E; A53E: 0.25±0.08, E46K: 0.34±0.10; p<0.05). A53E-α-synuclein also had a faster fluorescence recovery (tau) following photobleaching as compared to both WT-α-synuclein and E46K-α-synuclein (Figure 2F; A53E: 0.9 min±0.3; WT: 2.1 min±1.5; E46K: 2.7 min±1.2; p<0.005). This is consistent with previous *in vitro* data suggesting that A53E-α-synuclein forms smaller oligomers compared to WT-α- synuclein^19,21^. E46K-α-synuclein showed the lowest mobility with a higher immobile fraction relative to WT (0.24±0.06; p<0.001). These differences indicate that disease-causing mutations influence mobility differentially *in vivo*, with E46K decreasing mobility compared to A53E, possibly via the formation of distinct types of aggregates at this developmental stage.

**Figure Two.**
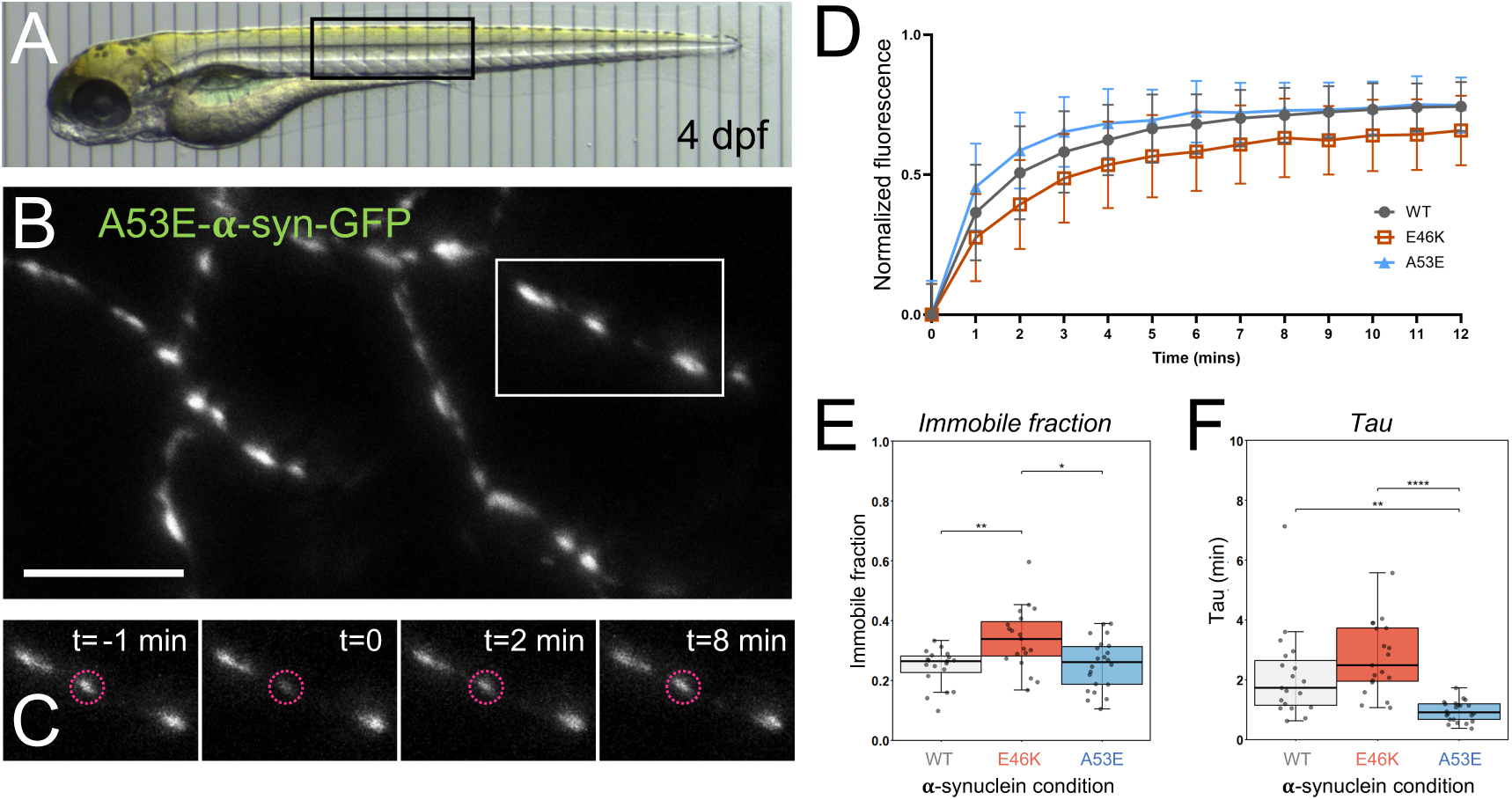
A53E-α-synuclein is more mobile than WT- and E46K-α-synuclein at 4 days. A) *In vivo* transmitted light image of zebrafish larva at 4 days post fertilization (dpf) with box indicating general region targeted for FRAP. B) Representative section of 4 dpf motor axon expressing A53E-α-synuclein-GFP, used for FRAP experiment. Box indicates region containing synaptic terminal targeted for photobleaching enlarged in C. Scale bar is 10 μm. C) Representative time series showing fluorescence recovery of a photobleached terminal before (t=-1 min) and during the course of one FRAP experiment. Dashed circles indicate the experimental terminal photobleached just before t=0, and the following images show the recovery of unbleached α-synuclein-GFP from neighboring terminals at 2 and 8 min. Panels in B-C are maximum intensity projections with the following depths: B) 7.7 μm; C) 5.6 μm. D) Normalized fluorescence recovery curves showing the average fluorescence intensity over time following photobleaching for all α-synuclein conditions (WT: dark gray line/circles, n=50; E46K: orange line/squares, n=52; A53E: blue line/triangles, n=51. E) Quantification of the amount of α-synuclein-GFP that does not recover at 4 dpf, as measured by immobile fraction (IF; WT: 0.24±0.06, n=50 terminals from 19 neurons across 16 larvae; E46K: 0.34±0.10, n=52 terminals from 19 neurons across 18 larvae; A53E: 0.25±0.08, n=51 terminals from 22 neurons across 19 larvae). IF differed significantly among conditions (Kruskal-Wallis, p=0 .004), with Dunn’s post-hoc revealing that E46K IF was significantly higher than both WT (p=0.007) and A53E (p=0.02). F) Quantification of α-synuclein mobility at 4 dpf as measured by tau (WT: 2.1±1.5 min; E46K: 2.7±1.2 min; A53E: 0.9±0.3 min). Tau differed significantly among conditions (Kruskal-Wallis, p=0.003), with Dunn’s post-hoc revealing that A53E tau was significantly lower than WT (p=0.002) and E46K (p<0.0001). For E and F, boxes represent the interquartile range, line indicates median, and whiskers indicate range, with individual neuron measurements overlaid. Each dot overlaid on the graph represents the average of several individual FRAP experiments completed at different synaptic terminals within one neuron’s axonal arbor. Statistical significance indicated by * p<0.05, ** p<0.01, **** p <0.0001.

### WT- and E46K-α-synuclein mobility progressively decreases over time, while A53E-α- synuclein remains relatively mobile

We next tested whether the distinct mobilities we measured at 4 dpf reflect stable differences among α-synuclein forms, or if they could potentially change over time, revealing progressive changes in α-synuclein aggregation state. Our *in vivo* model allows us to image the protein at multiple timepoints, thus we performed *in vivo* FRAP experiments at 2, 4, and 6 dpf in each condition (Figure 3). Across these developmental times, both WT- and E46K-α-synuclein mobility progressively decreased, as measured by an increasing immobile fraction from 2 to 6 dpf (Figure 3A-B; WT IF at 2 dpf: 0.23±0.12; WT 4 dpf: 0.24±0.06; WT 6 dpf: 0.44±0.13; p≤0.0009. E46K IF at 2 dpf: 0.30±0.09; E46K 4 dpf: 0.33±0.10; E46K 6 dpf: 0.53±0.15; p<0.005). Additionally, E46K recovery tau increased significantly over time (Figure 3C-D; 2 dpf: 1.5±0.82 min; 4 dpf: 2.7±1.2 min; 6 dpf: 4.9±2.3 min; p<0.04). This pattern suggests that the WT- and E46K- forms α-synuclein protein become increasingly more aggregated over days *in vivo*, as expected for these conditions. In contrast, A53E-α-synuclein surprisingly did not show the same progressive decline in mobility. Its IF remained stable across 2, 4, and 6 dpf (Figure 3A, right column; A53E IF at 2 dpf: 0.22±0.10; 4 dpf: 0.25±0.08; 6 dpf: 0.25±0.08); however, its mobility as measured by tau decreased slightly from 4 dpf to 6 dpf (Figure 3C, right panel; A53E tau at 2 dpf: 1.4±0.59, at 4 dpf: 0.95±0.35, at 6 dpf: 2.0±1.1 p<0.01). At later time points, as the mobility of E46K- and WT-α-synuclein progressively decreased, A53E-α- synuclein had significantly lower immobile fraction (Figure 3B) and/or tau values (Figure 3D) than both WT- and E46K-α-synuclein, suggesting that the A53E human disease mutation allows the α-synuclein protein to remain in a more mobile state for a longer period *in vivo*.

**Figure Three.**
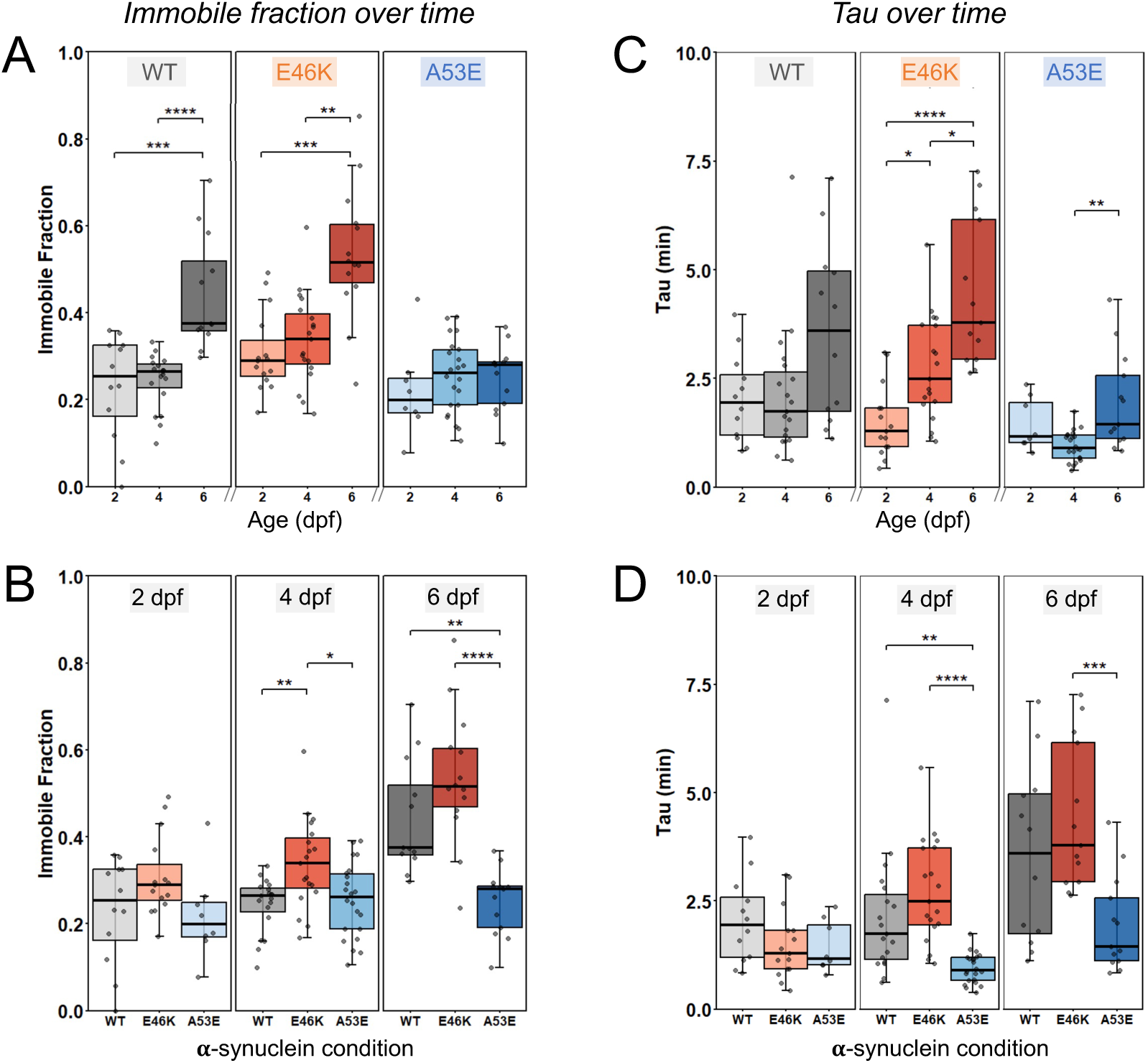
Wild-type- and E46K-α-synuclein aggregation increases over time *in vivo*, but A53E-α-synuclein remains mobile until at least 6 days. A) *In vivo* FRAP experiments quantified immobile fraction measured at 2, 4 and 6 days post fertilization (dpf), grouped by condition (WT IF 2 dpf: 0.23±0.11, n=32 terminals from 16 neurons across 9 larvae; WT IF 4 dpf: 0.24±0.06, n=50 terminals from 19 neurons across 16 larvae; WT IF 6 dpf: 0.44±0.13, n=30 terminals from 12 neurons across 9 larvae; Kruskal Wallis, p<0.0001; Dunn’s multiple comparison 2 dpf to 6 dpf p=0.0009; 4 dpf to 6 dpf p<0.0001. E46K IF 2 dpf: 0.30±0.09, n=37 terminals from 15 neurons across 12 larvae; E46K IF 4 dpf: 0.33±0.10, n=52 terminals from 19 neurons across 18 larvae; E46K IF 6 dpf: 0.53±0.15, n=40 terminals from 13 neurons across 10 larvae; Kruskal Wallis, p=0.0001; Dunn’s multiple comparison 2 dpf to 6 dpf p=0.0002; 4 dpf to 6 dpf p=0.002. A53E IF 2 dpf: 0.22±0.10, n=21 terminals from 8 neurons across 7 larvae; A53E IF 4 dpf: 0.25±0.08, n=51 terminals from 22 neurons across 19 larvae; A53E IF 6 dpf: 0.25±0.08, n=36 terminals from 13 neurons across 11 larvae. Kruskal Wallis, p=0.4). B) Same immobile fraction values as in A, now grouped by age. Center panel data (4 dpf) is duplicated from Figure 2 for direct comparison with 2 and 6 dpf. Significant differences between conditions seen at 4 and 6 dpf, with E46K exhibiting a greater immobile fraction than WT (p=0.007) and A53E (p=0.02) at 4 dpf, and A53E exhibiting a smaller immobile fraction than both WT (p=0.003), and E46K (p<0.0001) at 6 dpf. C) Tau measured at 2, 4, and 6 dpf, grouped by condition, same sample sizes as A. WT-α-synuclein tau at 2 dpf: 2.0±1.0 min; WT tau 4 dpf: 2.1±1.4 min; WT tau 6 dpf: 3.5±2.1 min; Kruskal Wallis, p=0.1. E46K-α-synuclein tau at 2 dpf: 1.5±0.82 min; E46K tau 4 dpf: 2.7±1.2 min; E46K tau 6 dpf: 4.9±2.3 min. 2 dpf to 6 dpf p<0.0001; 4 dpf to 6 dpf p=0.04. A53E-α-synuclein tau at 2 dpf: 1.4±0.59 min; A53E tau 4 dpf: 0.95±0.35 min; A53E tau 6 dpf: 2.0±1.1 min. 4 dpf to 6 dpf p=0.004. D) Same tau values as in C, now grouped by age. Significant differences between conditions are seen at 4 and 6 days, with A53E having a smaller tau than WT and E46K at these ages (4 dpf: A53E to WT p=0.002; A53E to E46K p<0.0001. 6 dpf: A53E to E46K p=0.0009). Center panel data (4 dpf) is duplicated from Figure 2 for direct comparison with 2 and 6 dpf. Boxes in A-D represent the interquartile range, line indicates median, and whiskers indicate range. Each dot overlaid on the graph represents the average of several individual FRAP experiments completed at different synaptic terminals within one neuron’s axonal arbor. Statistical significance indicated by * p<0.05, ** p<0.01, *** p <0.001, and **** p<0.0001.

These data reveal patterns about the progression of the α-synuclein population over time *in vivo*; we next wanted to test how α-synuclein protein within one individual neuron’s cytoplasm behaves over time, and whether the slowing of WT- and E46K-α-synuclein occurs within individual neurons or simply reflects heterogeneous progression across the population. To do this, we developed new techniques to use longitudinal *in vivo* FRAP within individual neurons over time. A given neuron within a zebrafish larva was imaged first at 2 dpf, then the same neuron was located and imaged again at 4 dpf and 6 dpf (Figure 4A-B). At each time point, we measured mobility in multiple presynaptic terminals within the same neuron. While it was not possible to identify the exact same terminals due to developmental changes, we performed FRAP in up to five separate terminals within each neuron at each time point, and calculated the average value across those terminals over time for each individual neuron. Using this approach, we found that WT- and E46K-α-synuclein mobility decreased over time within a given neuron, demonstrated by an increased immobile fraction, consistent with the population data (Figure 4C; WT and E46K; Supplemental Figure 2). This change in mobility over time was then represented as the slope of the increasing immobile fraction from 2 to 6 dpf, and graphed as the propensity to aggregate (Figure 4D; WT 0.35±0.06, E46K 0.24±0.07). By contrast, in neurons expressing A53E-α-synuclein, mobility did not increase over time but stayed relatively constant, revealing a significantly lower propensity to aggregate (Figure 4C lower panel and 4D; A53E 0.05±0.04; p≤0.003). This suggests that even within individual neurons *in vivo*, the cytoplasmic population of WT- and E46K-α-synuclein protein progressed in its aggregation state, while the A53E mutation rendered α-synuclein less likely to aggregate.

**Figure Four.**
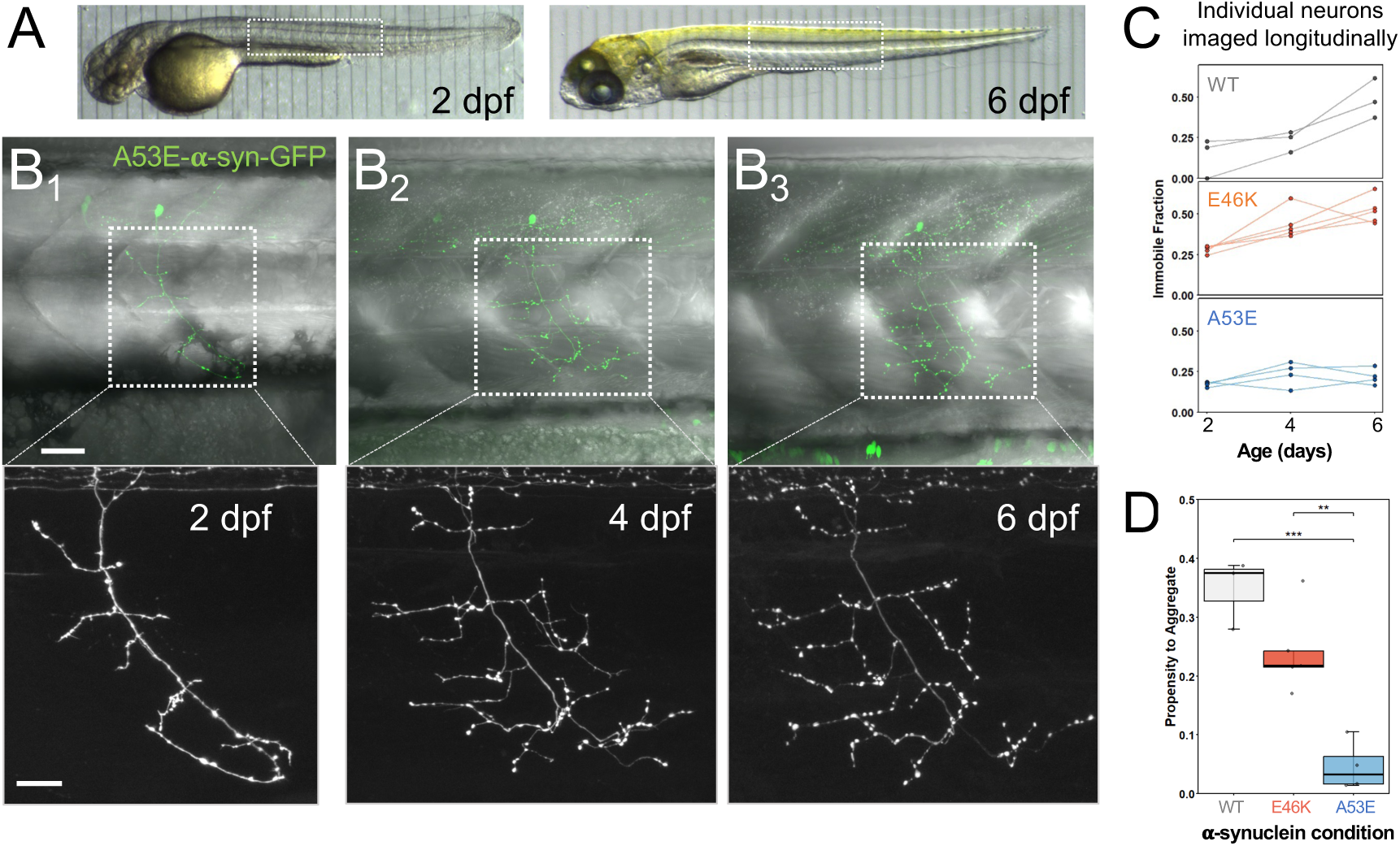
Within individual neurons over time *in vivo*, A53E-α-synuclein remains mobile, while both wild-type- and E46K-α-synuclein become less mobile. A) *In vivo* transmitted light images of larval zebrafish at 2 dpf (left) and 6 dpf (right). Dotted boxes represent general region targeted for FRAP. B) Longitudinal imaging of the same α-synuclein-expressing motor neuron at 2 dpf (B1), 4 dpf (B2), and 6 dpf (B2). Upper panels show GFP fluorescence channel overlaid with transmitted light to show trunk morphology and *in vivo* context. Dotted boxes in B indicate regions magnified in lower panels, which show distinct axon development over time. Scale bar 40 μm in upper images, 20 μm in lower. Panels in B are maximum intensity projections of Z-stacks with the following depths: 2 dpf/B1) 53.2 μm; 4 dpf/B2) 93.1 μm; 6 dpf/B3) 96.6 μm. C) Immobile fraction (IF) of α-synuclein measured in the same individual neurons longitudinally at 2, 4, and 6 dpf for WT (top, grey), E46K (middle, orange), and A53E (bottom blue). Each dot represents the average IF within one individual neuron at that time point (averaged across up to five synaptic terminals), and lines connect measurements from one same neuron across time. Individual values are provided in Supplemental Figure 2. D) Propensity to aggregate as measured by the slope of the lines (from 2 to 6 dpf) represented in (C). For WT-α-synuclein-expressing neurons, propensity is 0.35±0.06; E46K is 0.24±0.07; A53E is 0.05±0.04. A one-way ANOVA revealed A53E has a significantly lower propensity to aggregate than both WT and E46K (p=0.0003; post-hoc Tukey test A53E to WT p=0.0003, A53E to E46K p=0.003, WT to E46K p=0.09). Boxes represent the interquartile range, line indicates median, and whiskers indicate range. Each dot represents the propensity for α-synuclein to aggregate within one individual neuron, i.e., the slope of one line from 2-6 dpf in C). ** p<0.01, and *** p<0.001.

### A53E-α-synuclein lacks detectable phosphorylation at serine-129

Since the A53E mutation appears to influence α-synuclein mobility *in vivo*, we next tested how this mutation correlates with disease-relevant post-translational modifications. One key feature of α-synuclein pathology is phosphorylation at serine-129 (pS129), which is found in over 90% of α-synuclein within Lewy bodies^44^. Phosphorylation at this site has been studied extensively as a possible driver of aggregation; however, in at least some studies, phosphorylation at S129 alone does not cause α-synuclein to aggregate^27,45,46^. This raises the possibility that phosphorylation may require a more complex set of conditions in order to drive aggregation, and/or may be a late-stage consequence of α-synuclein aggregation rather than the driving force. Additionally, some studies suggest that pS129 may increase with PD progression, as it correlates with disease severity^47^, further supporting the hypothesis that pS129 could serve as a marker of disease progression rather than an early driver of disease. Currently, it is not known how patterns of S129 phosphorylation relate to human mutations in α-synuclein in the living nervous system. Examining pS129 in the context of these mutations *in vivo* could provide insights into its relationship with aggregate size and state.

We have previously shown that WT-α-synuclein is phosphorylated at S129 by endogenous Polo-like kinases in zebrafish, as it is in mammalian cells^27^. To test whether E46K- and A53E-α- synuclein are phosphorylated at similar levels, we performed whole-mount immunohistochemical staining using a human α-synuclein antibody that specifically recognizes α-synuclein phosphorylated at serine-129. As described in Weston et al.^27^, axonal arbors typically showed either full staining throughout the majority of synaptic terminals, or none. Surprisingly, we found that S129 phosphorylation appeared to be abolished in A53E-α-synuclein at 4 dpf (Figure 5A-B). While phosphorylation was detected in both WT-and E46K-α-synuclein-expressing conditions, no phosphorylation was detected in any A53E motor neuron, indicating that the A53E mutation disrupts and/or delays phosphorylation at S129 *in vivo* (Figure 5B; WT: 32%, n=57; E46K: 7%, n=41; A53E: 0%, n=45).

**Figure Five.**
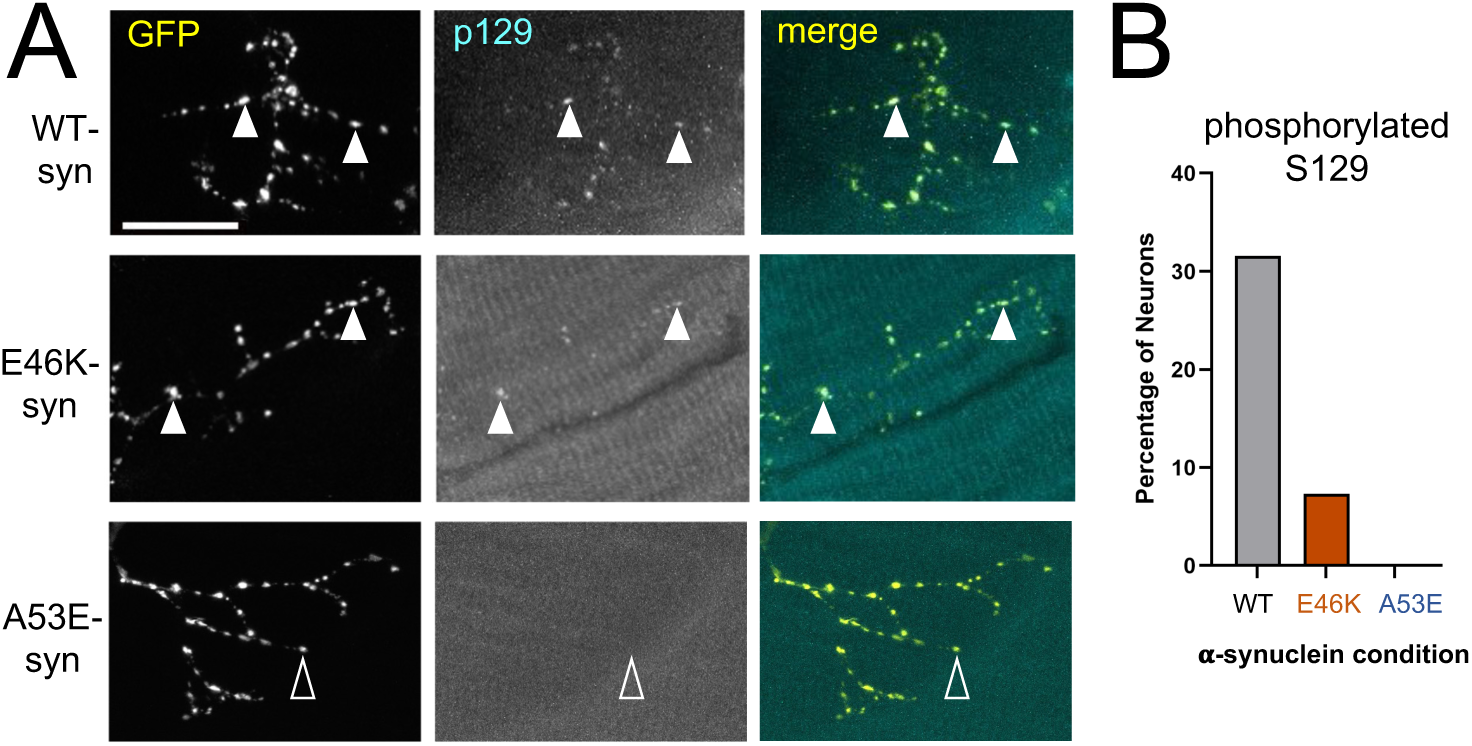
A53E mutation to α-synuclein abolishes phosphorylation of serine-129. A) Whole-mount immunohistochemical staining using a human anti-phospho-α-synuclein antibody that specifically recognizes α-synuclein phosphorylated at serine-129 in 4 dpf larvae expressing WT- (top) and E46K- (middle), or A53E-α-synuclein (bottom). Left panels show α-synuclein-GFP channel; middle panels show anti-phospho-S129; right panels show merge. Scalebar is 20 um. Filled white arrowheads show examples of positive colocalization, while empty arrowheads highlight no staining. Panels in A are maximum intensity projections of Z-stacks with the following depths: WT 17.85 μm; E46K 14.79 μm; A53E 16.32 μm B) Graph representing the percentage of neurons with positive staining for phosphorylation at S129 (WT: 32%, n=57; E46K: 7%, n=41; A53E: 0%, n=45; see Methods for criteria).

### Mimicking phosphorylation at S129 is sufficient to decrease A53E-α-synuclein mobility

The absence of detectable pS129 in A53E-expressing neurons raised the possibility that the phosphorylation state of α-synuclein could contribute to the distinct mobility behavior found with the A53E mutation. Because A53E-α-synuclein remained relatively mobile and did not show the progressive decrease in mobility we observed for WT- or E46K-α-synuclein, we asked whether mimicking phosphorylation at S129 alongside the A53E-α-synuclein would be sufficient to alter this behavior. To answer this question, we created a phosphomimetic form of A53E-α- synuclein where serine-129 is replaced with aspartate to mimic the negative charge of phosphorylation at this site (A53E+S129D; Figure 6A). We then compared the mobility of A53E+S129D to the other three forms of α-synuclein at 2 dpf, 4 dpf, and 6 dpf, both at the population level (Figure 6B) and longitudinally in individual neurons (Figure 6C-D), to test whether this forced phosphorylation mimic at S129 would be sufficient to shift α-synuclein into a less mobile state. Indeed, we found that the IF of A53E+S129D-α-synuclein increased over time, in a pattern similar to WT and E46K (Figure 6B; A53E+S129D IF at 2 dpf: 0.21±0.08; 4 dpf: 0.31±0.05; 6 dpf: 0.37±0.11; p<0.05). This suggests that mimicking S129 phosphorylation returned A53E-α-synuclein to a less mobile, more aggregated state. Similarly, when we performed FRAP experiments in single neurons over time, we measured a progressive decrease in mobility of A53E+S129D-α-synuclein, as measured by a higher immobile fraction over time (Figure 6C-D and Supplemental Figure 2), the same pattern seen in both WT- and E46K-α- synuclein conditions. These results suggest that the forced phosphorylation at S129D was sufficient to increase the propensity of A53E-α-synuclein to aggregate *in vivo*.

**Figure Six.**
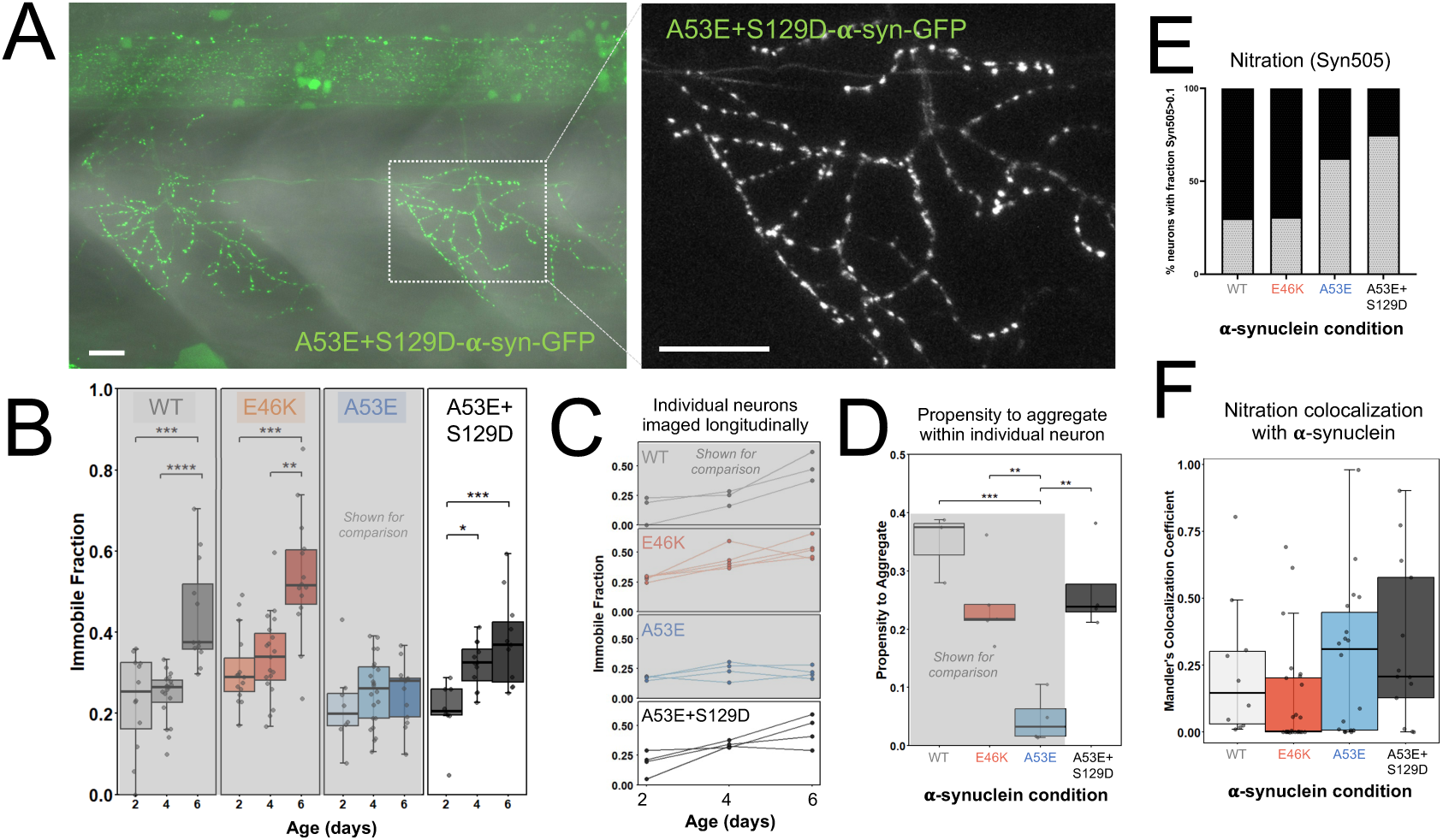
Genetic modification to mimic phosphorylation in A53E-α-synuclein is sufficient to decrease its mobility *in vivo*. In order to compare A53E+S129D directly to the other conditions, some data in panels B-D (grayed out) are repeated from Figures 3 and 4. A) *In vivo* image of A53E+S129D-expressing 4 dpf zebrafish with GFP fluorescence channel overlaid over transmitted light to show trunk morphology and *in vivo* context (left). Dotted box shows region enlarged in panel at right, highlighting punctate axon morphology. Both panels show a maximum intensity projection of 60.9 μm. Scale bar is 20 μm. B) A53E+S129D-α-synuclein immobile fraction increases over time (Kruskal Wallis p=0.005) from 2 dpf to 4 dpf (p=0.04, Dunn’s multiple comparison) and from 2 to 6 dpf (p=0.004; IF at 2 dpf: 0.21±0.08, n=16 terminals from 7 neurons across 6 larvae; 4 dpf: 0.32±0.06, n=28 terminals from 11 neurons across 9 larvae; 6 dpf: 0.37±0.12, n=27 terminals from 10 neurons across 7 larvae). Each dot overlaid represents the average of several individual FRAP experiments done at multiple synaptic terminals within one neuron. Leftmost three panels are repeated from Figure 3 for direct comparison with A53E+S129D condition. C) Alpha-synuclein immobile fraction (IF) measurements from the same individual neurons imaged longitudinally at 2, 4, and 6 dpf with A53E+S129D condition included (bottom panel) for direct comparison. Each dot represents the average IF within one individual neuron at that time point, and lines connect measurements from the same neuron across time. Individual values are provided in Supplemental Figure 2. Upper three panels are repeated from Figure 4 for direct comparison with A53E+S129D condition. D) Propensity to aggregate as measured by the average slope of the lines in C for each condition. Each dot represents one neuron imaged longitudinally at 2, 4, and 6 dpf. For A53E+S129D-α-synuclein-expressing neurons, value is 0.27+/-0.08. Leftmost three bars are repeated from Figure 4 for direct comparison with A53E+S129D condition. ANOVA overall p=0.0003; A53E to A53E+S129D, Tukey p=0.002. E) Graph representing the percentage of neurons with nitration colocalization with α-synuclein at levels greater than 0.1 (WT: 30% n=10; E46K: 31% n=25; A53E: 63% n=18; A53E+S129D: 75% n=13). F) A Mander’s colocalization analysis revealed quantified levels of nitrated α-synuclein for WT-α-synuclein (0.22±0.25; n=10 neurons from 3 larvae), E46K-α-synuclein (0.12±0.20; n=25 neurons from 11 larvae), A53E-α-synuclein (0.27±0.28; n=18 neurons from 10 larvae), and A53E+S129D-α-synuclein (0.32±0.30; n=13 neurons from 6 larvae); Kruskal Wallis, p=0.02.

Alpha-synuclein aggregation has been linked to impaired mitochondrial function and the promotion of oxidative stress^48,49^. As such, we wished to test how each form of α-synuclein was nitrated, as a potential indicator of oxidative stress and aggregation. We specifically wanted to test whether A53E-α-synuclein, which is known to cause a more virulent form of PD but was found to be the most mobile in our experiments, was more or less nitrated than the other forms of α-synuclein. We measured the nitration state of each of the α-synuclein conditions using an antibody that recognizes nitrated α-synuclein and specifically binds misfolded forms of the protein^50^. As expected, we found some amount of nitration in each of our four conditions (WT-α- synuclein, E46K-α-synuclein, A53E-α-synuclein, and A53E+S129D-α-synuclein; Figure 6E-F). Interestingly however, unlike the clear prevention of S129 phosphorylation, we found robust nitration of A53E-α-synuclein. Forcing the phospho-mimic at S129 did not ameliorate these nitration levels in A53E-α-synuclein, but appeared to slightly increase them. E46K-α-synuclein, on the other hand, the form that showed the lowest mobility (consistent with higher aggregation), appeared to have a lower amount of nitration. These data suggest that the A53E mutation maintains or may increase nitration of α-synuclein, irrespective of S129 phosphorylation state.

## Discussion

Rare, disease-associated α-synuclein mutations A53E and E46K lead to diverging PD outcomes, but their aggregation dynamics have not yet been studied or compared *in vivo*. Our results demonstrate that these mutations cause distinct effects on α-synuclein dynamics *in vivo* and that these effects can be measured over time, both at the population level and within individual neurons. Using *in vivo* FRAP to measure the mobility of WT-, E46K-, and A53E-α-synuclein in presynaptic terminals, we found that A53E-α-synuclein was significantly more mobile than both WT- and E46K-α-synuclein. A53E-α-synuclein maintains this high mobility over four days *in vivo,* while both WT- and E46K-α-synuclein become progressively more immobile during this period. These results suggest that A53E-α-synuclein aggregates less or at a slower rate than other forms of α-synuclein. We developed methods to measure this pattern in single neurons across three time points, offering a novel way to investigate cytoplasmic protein dynamics and the early stages of protein aggregation over time in the living nervous system. Our longitudinal approach within synaptic terminals of individual neurons *in vivo* is particularly valuable because it reflects real changes occurring at the cellular level within the context of the living nervous system.

Our findings indicate that a single residue alteration can shift α-synuclein into distinct aggregation states *in vivo*. Consistent with other findings, our data support the idea that the A53E mutation in α-synuclein may favor formation of a smaller, more mobile oligomeric species, while inhibiting further aggregation into larger, less mobile protofibril or fibril forms^18,19,21,51,52^. Such smaller oligomeric species would be expected to have a faster recovery tau in FRAP experiments and possibly a decreased immobile fraction^37^, consistent with what we measured for A53E. The cause for this shift in the steady-state towards more mobile oligomeric species is unclear, but could be due to the nature of the negatively charged glutamate at this position, since Maji and colleagues have studied several different uncharged amino-acids at this position, all of which promote larger fibrillar aggregates more than A53E^52^. The glutamate residue at this site may thus interfere with the formation of fibrils, potentially via electrostatic forces.

We also observed that A53E-α-synuclein lacks detectable phosphorylation at the S129 residue, which may contribute mechanistically to its distinctly high mobility. S129 phosphorylation (pS129) is one of the most prominent post-translational modifications associated with pathological human α-synuclein in PD, yet its relationship with and impact on aggregation remain unclear. Phosphorylation has been associated with the aggregation of key proteins in other neurodegenerative diseases as well, such as Aß in Alzheimer’s disease^53,54^. Although pS129 is highly enriched in Lewy bodies, several previous studies have shown that phosphorylation at this site alone is not sufficient to drive α-synuclein aggregation^27,45,46^. Our lab has previously demonstrated that WT-α-synuclein can be phosphorylated by endogenous Polo-like kinases in zebrafish^27^, as in mammalian cells. Here, as expected, this endogenous phosphorylation was detected in neurons expressing both WT- and E46K-α-synuclein; however, it was abolished in A53E-α-synuclein expressing neurons. This is consistent with previous *in vitro* studies reporting reduced S129 phosphorylation of A53E-α-synuclein in human neuroglioma cells^21^. Quantification of phosphorylation levels in human tissue derived from A53E-induced PD patients is an important question for future studies, although challenging, given the rare nature of this variant (currently only 3-4 known families). Our vertebrate *in vivo* approach provides insights into the relationship between phosphorylation and A53E aggregation, a dynamic that remains poorly understood in human A53E-associated pathology.

The unique absence of detectable S129 phosphorylation in A53E-expressing neurons reveals a potential mechanism for the distinct mobility trajectory of this mutation. To test this directly, we designed a mutant form of α-synuclein that mimics its phosphorylation at S129 (A53E+S129D). Surprisingly, we found that in this condition, α-synuclein showed a progressive decrease in mobility, more similar to the WT- and E46K-forms of the protein. This suggests that S129 phosphorylation, although it may not drive aggregation on its own^27^, can impact the aggregation tendency of the protein in the context of other specific mutations. One possibility is that the lack of pS129 in A53E-α-synuclein contributes to its persistence in a more mobile state, while the presence of a negative charge at this position (caused by either phosphorylation or the mimic aspartate) promotes a conformational change that favors progression toward larger aggregates like fibrils. Taken together, our results show that the presence of detectable pS129 can differ among α-synuclein conditions *in vivo,* raising the possibility that the phosphorylation state of S129 may contribute to the different α-synuclein aggregation trajectories, even if it does not act as the sole driver of aggregation.

Our nitration data indicated that the relationship between α-synuclein aggregation and nitration state is more complex than a direct association with aggregation. Notably, both A53E- and A53E+S129D-α-synuclein appear to exhibit high levels of nitration at 4 dpf, despite their differing mobility patterns. One possible explanation is that A53E promotes the formation of oligomeric species that contribute to toxicity and are particularly susceptible to oxidative stress. The S129D substitution may alter A53E-α-synuclein aggregation dynamics by progressively shifting the protein toward less mobile species; however, nitration could occur before this shift and persist through the aggregation process. This could help explain the somewhat elevated nitration that we observed in A53E+S129D-α-synuclein relative to E46K-α-synuclein, despite their similar mobility measurements. In addition, nitration may progress at different rates for different forms of α-synuclein and could be measured at multiple time points.

Perhaps the most notable aspect of the high mobility observed in A53E-α-synuclein is that it is coupled with particularly severe disease and mortality outcomes in patients who carry this mutation. One possible explanation is that oligomeric species are particularly toxic^5,22^, and that cells may respond by modifying α-synuclein through phosphorylation at S129, which could promote the sequestration of these small oligomers into larger inclusions or toward degradation, to minimize toxic effects^21,55^. The A53E substitution in α-synuclein, however, could impinge upon the cell’s ability to phosphorylate the protein at this site, maintaining the protein in a more mobile, oligomeric state for an extended period. This could cause more harm to the cell while the oligomeric species persist, driving more severe disease outcomes.

Studying familial α-synuclein mutations offers a direct way to determine how specific changes in the protein sequence alter its behavior *in vivo* and contribute to disease. Because these mutations alter α-synuclein aggregation and mobility, they provide a means to identify what pathways may contribute to the formation of Lewy bodies and, ultimately, neuronal dysfunction and neurodegeneration. Importantly, the mechanisms revealed by familial mutations are also relevant to sporadic PD, where changes in α-synuclein aggregation occur in the absence of a pathogenic *SNCA* mutation. Therefore, understanding how individual α-synuclein mutations alter protein dynamics in living neurons provides a valuable framework for understanding the cellular mechanisms that regulate α-synuclein aggregation more broadly. Defining these mechanism*s in vivo* is an important step towards connecting changes in α-synuclein to neuronal dysfunction and the clinical progression of PD, while providing insights into pathological protein aggregation that may be broadly shared across neurodegenerative diseases.

## Methods

### Generation of DNA constructs

DNA sequences of 1) human wild-type (WT)-α-synuclein (gift from P. McLean, Mayo Clinic); 2) A53E-α-synuclein; 3) E46K-α-synuclein; 4) A53E+S129D-α-synuclein; and 5) EGFP were flanked with attB1 and attB2 sites for Multisite Gateway Cloning®. Each modified version of α- synuclein was designed in SnapGene® software (from Dotmatics, available at snapgene.com) and DNA fragments were synthesized by Integrated DNA Technologies (Coralville, IA). PCR products were then recombined with Gateway pDONR221, creating middle entry clones. Entry clones were transformed into 5-alpha *E. coli* (New England Biolabs, Ipswich, MA) and DNA was extracted and purified through the QIAprep Spin Miniprep Kit (QIAGEN, Hilden, Germany). Middle elements were recombined with pDestTol2CG (pDESTTol2pA with cmlc2:egfp transgenesis marker; Kwan et al., 2007), a NeuroD promoter as the 5’ element (p5E containing a ∼5 kb segment of promoter/enhancer sequence upstream of the neuroD gene; gift from T. Nicolson), and a 3’ EGFP. The EGFP construct includes the 3’ SV40 late poly-A element (p3E-polyA^56^, in place of the EGFP p3E (See Table 1). Final expression clones were amplified by transformation into 10-beta *E. coli* (New England Biolabs, Ipswich, MA), and plasmid DNA was purified by miniprep, and sequences were confirmed by whole plasmid sequencing (Plasmidsaurus, San Francisco, CA).

**Table 1.** DNA Plasmids used.

| Abbreviation | Plasmid Name | 5' Element | Middle Element | 3' Element |
| --- | --- | --- | --- | --- |
| WT-α-synuclein | pEXP_NeuroD_$\alpha$_synuclein_EGFP | NeuroD Promoter | WT-α-synuclein | EGFP |
| A53E-α-synuclein | pEXP_NeuroD_A53E_$\alpha$_synuclein_EGFP | NeuroD Promoter | A53E-α-synuclein | EGFP |
| E46K-α-synuclein | pEXP_NeuroD_E46K_$\alpha$_synuclein_EGFP | NeuroD Promoter | E46K-α-synuclein | EGFP |
| A53E+S129D-α-synuclein | pEXP_NeuroD_A53E+S129D_$\alpha$_synuclein_EGFP | NeuroD Promoter | A53E+S129D-α-synuclein | EGFP |
| GFP | pEXP_NeuroD_EGFP_pA | NeuroD Promoter | EGFP | polyA |

### Zebrafish maintenance and expression of DNA constructs

Zebrafish protocols were approved by Lewis & Clark College’s Institutional Animal Care and Use Committee (IACUC). Wild-type AB, TL, and AB/TL lines of adult zebrafish were maintained and used for matings; all experiments were conducted in AB/TL embryos. Adult zebrafish were reared in a controlled, circulating, and monitored tank facility with reverse osmosis water at 27℃ and a 12/12 hour light/dark cycle (Aquaneering Inc.; San Diego, CA). Adult zebrafish were fed dry food (Zebrafeed by Sparos I&D; Olhão, Portugal) and brine shrimp (*Artemia*; Brine Shrimp Direct; Ogden, UT). After mating adult zebrafish, fertilized eggs at the one-cell stage were microinjected with DNA plasmids within 45 minutes of fertilization under a dissecting microscope (Olympus SZ61, Tokyo, Japan). DNA injection mix consisted of the DNA plasmid diluted to 10-13 ng/µL using 0.1M KCl (Alfa Aesar, Ward Hill, MA) and 1% phenol red dye (Sigma Aldrich, St. Louis, MO) to allow for visualization during injections. Injection mix was loaded into a small pulled (P-97, Sutter Instrument Co., Novato, CA) glass capillary needle (WPI TW100F-4 World Precision Instruments, Sarasota, FL), and injected using a nitrogen gas-powered microinjection system (World Precision Instruments; Sarasota, FL, USA). The amount of DNA ejected was calibrated to a 200 μm bolus size, which corresponded to about 0.042 ng of DNA released per embryo. Injected embryos were stored in E3 embryo medium^57^ at 28℃ (Benchmark Scientific; Sayreville, New Jersey). At 24 h post fertilization (hpf), embryos were transferred to E3 with 0.2 mM phenylthiourea (PTU; Alfa Aesar, Ward Hill, MA) to inhibit development of skin pigment.

### Live Imaging and FRAP

Imaging of live larvae was performed similarly to previously described^27^. In preparation for live imaging, zebrafish aged 2-6 dpf were anesthetized in a solution of ∼0.2 mM Syncaine MS-222 (tricaine methanesulfonate; Syndel, Ferndale, Washington). Larvae were mounted onto glass slides in 1% low-melt agarose (AMRESCO Agarose SFR™; AMRESCO, Solon, OH) and imaged using an upright confocal microscope (LSM 710 Carl Zeiss, Oberkochen, Germany) under a 20X water immersion (1.0NA) objective. Experiments conducted in motor neurons were imaged at 8X zoom using ZEN black software (Zeiss ZEN Microscopy Software, Jena, Germany). Alpha-synuclein was visualized using a 488 nm argon laser (MBS 488/561) that excited GFP with a collection range of 493-598 nm (laser power=0.3-2, gain=600-700).

Fluorescence Recovery After Photobleaching (FRAP) experiments were performed to measure protein mobility in each condition. Motor axons projecting ventrally were located and distinguished using their branching patterns and characteristic large presynaptic terminals^58^. Experimental terminals for FRAP were located sufficiently far from branch points and the branch terminus, to maintain consistency in the amount of protein available to for diffusion. The terminal’s brightest point was centered in the Z-stack and an ROI was drawn around the experimental terminal selected for bleaching. A 9-slice Z-stack (0.7 μm interval) through the terminal was acquired at one-minute intervals for a total of 14 minutes. Just before the second image was collected (timepoint zero), the experimental terminal was bleached with a 100% 488 nm argon laser, a measure that was found to bleach terminals to at least 40% of their baseline fluorescence intensity. During and following image collection, the experimental terminal was assessed for drift. If the brightest plane moved out of the imaged volume during the course of the 14-minute experiment, the experiment was deemed unusable.

### FRAP Analysis

Analysis of FRAP image series was performed using Fiji^59^. Maximum intensity projection from each single bleached terminal was condensed into a time series and X/Y drift-corrected using the StackReg plugin (Biomedical Imaging Group, Swiss Federal Institute of Technology, Lausanne, Switzerland). To quantify intensity over time within the experimental terminal, a semi-automated method was first used to generate a region of interest (ROI): the first time point image was duplicated, a Gaussian blur (σ = 1.00) was applied and the threshold was manually set to encompass approximately 95% of the fluorescence. The resulting ROI was subsequently applied to the registered, unmodified time series and average fluorescence intensity within the ROI was measured for all 14 time points. Raw fluorescence intensity was then normalized to the baseline fluorescence of the terminal pre-bleach, to determine the percent fluorescence intensity throughout the experiment. Values were entered into Prism (GraphPad, La Jolla, CA) and fitted to a one-phase association exponential regression model. Time constant (tau), representing the time at which fluorescence had recovered to (1-1/e), or ∼63% of the final fluorescence value, was measured from the association curve. The immobile fraction (IF) of α-synuclein was calculated as follows, where initial value=1 and final value=plateau value obtained from Prism.

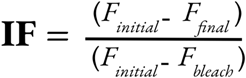

### Whole-mount immunohistochemistry

Immunohistochemistry was performed as previously described^27^. Each step was conducted at room temperature with light agitation unless otherwise noted. Four-day-old zebrafish larvae used for immunohistochemistry were fixed overnight in 4% paraformaldehyde (PFA; Electron Microscopy Sciences, Hatfield, PA) at 4°C. Then, they were washed in PBST (Phosphate Buffered Saline + 0.1% TritonX-100; AMRESCO, Solon, OH; Sigma Aldrich, St. Louis, MO) 3X for 15 minutes each, moved into PBS, and stored at 4°C for up to seven days. Larvae were incubated in acetone (EM Science, Gibbstown, NJ) for 15 minutes at −20°C, and washed with PBST once for 5 minutes and twice with PBS-DT (PBS, 1% BSA, 1% DMSO, 1% TritonX-100; Sigma Aldrich, St. Louis, MO) for 15 minutes. Larvae were then transferred to a blocking solution (PBS-DT, 5% NGS; Rockland Immunochemicals, Gilbertsville, PA and Sigma Aldrich, St. Louis, MO) for 90 minutes. Primary antibodies used included: 1) rabbit anti-phospho-S129 primary antibody (1:200; EP1536Y; Abcam, Cambridge, MA); and 2) mouse monoclonal anti-nitrated α-synuclein (1:100; nSyn505; Thermo Fisher Scientific, Waltham, MA). Each antibody was diluted in antibody buffer (20% blocking solution in PBS, 0.05% sodium azide; Sigma Aldrich, St. Louis, MO) and incubated for either 72 hours (anti-nitrated α-synuclein) or five days (anti-p-S129). The primary antibody was rinsed with PBST twice, followed by a PBST wash overnight. Secondary antibodies (diluted in antibody buffer and incubated for 2 hours) included: 1) anti-mouse Rhodamine Red-X-conjugated secondary antibody (1:500; Jackson ImmunoResearch, West Grove, PA) and 2) anti-rabbit Rhodamine Red-X-conjugated secondary antibody (1:400; Jackson ImmunoResearch, West Grove, PA). Finally, larvae were washed with PBST overnight, then prepared for imaging.

### Imaging of fixed larvae

Fixed larvae were mounted onto glass slides in Vectashield mounting medium (Vector Laboratories, Burlingame, CA) under a glass coverslip (VWR, Radnor, PA) supported by vacuum grease (Dow Corning Corp., Midland, MI) on each corner, and the sides were sealed with nail polish (essie, Montréal, Canada). Images of motor neurons projecting from the spinal cord were taken using either an upright or inverted Zeiss laser-scanning confocal microscope (LSM; Carl Zeiss LSM 710, Oberkochen, Germany) under a 40X oil (1.3 NA) objective. Motor neurons were identified based on their classic morphology (Myers et al., 1986). GFP was visualized using a 488 nm argon laser that excited GFP (MBS 488/561), with a collection range of 493-556 nm. Anti-pS129 or anti-nitrated α-synuclein signals were visualized using a 561 nm laser, with a collection range of 566-689 nm. Slides were stored in a dark environment at 4℃ for up to seven days prior to imaging. All confocal images (live and fixed) are maximum intensity projections with depths noted in figure legends. Whole fish brightfield images (Figures 1A, 2A, 4A) were taken using a Zeiss Discovery V8 dissection microscope (Oberkochen, Germany). Following GFP fluorescence image acquisition, identical linear adjustments were made to images within each figure for visual clarification.

### Immunohistochemistry Analysis

Synaptic terminals throughout the axonal arbor of each GFP-expressing neuron were imaged and examined for overlapping GFP and Rhodamine Red-X signal. Neurons were classified as positive if multiple synaptic terminals exhibited a Rhodamine Red-X signal above background on the brightest focal plane of a GFP terminal. Colocalization analysis of nitrated α-synuclein was performed at 4 dpf as previously described (Weston et. al., 2021), using maximum intensity projections with the JaCoP plugin^60^ in Fiji. Stacks were cropped in X, Y, and Z to include only relevant GFP fluorescence in motor neurons. Mander’s colocalization coefficients for GFP were calculated after setting intensity thresholds in each channel to one intensity level above the level where individual pixels in background areas were no longer detected. To control for the effect of thresholding, each image was independently thresholded and analyzed three separate times, and the mean Mander’s coefficient was used. A secondary quantification of nitration was done to address background noise and non-specific binding that could impact the colocalization analysis. The percentage of neurons with a Mander’s colocalization value of greater than 0.1 was calculated, a conservative cutoff for colocalization above background.

In order to quantify the punctate nature of GFP expression, a coefficient of variation analysis was used. Each axonal arbor, expressing either free GFP or WT-α-synuclein-GFP was roughly traced by hand in Fiji to create a region of interest (ROI), and the average fluorescence intensity and standard deviation within the ROI were calculated. A coefficient of variation was calculated (CV=standard deviation/average fluorescence intensity). A higher CV corresponded with higher variation in fluorescence intensity throughout the arbor and, therefore, increased punctate expression. A lower CV corresponded with a lower degree of variability and, therefore, more diffuse, even fluorescence.

### Presynaptic Terminal Quantification

To quantify the distance between and size of presynaptic terminals in motor neurons, Z-stacks were 3D-rendered in Imaris image analysis software (version 9.9.1, Oxford Instruments). The axonal arbors were manually traced using the Filaments function, and the presynaptic terminals were detected semi-automatically using the Spots function, with secondary visual confirmation. Spot detection parameters were established using WT-α-synuclein neurons, where the diameter and histogram thresholding were manually performed to set an estimated criteria range of intensity values for a presynaptic terminal. For each neuron, three to four segments of the axonal arbor at varying locations were identified, manually traced, and analyzed for terminals. To calculate the distance between presynaptic terminals, the length of each filament segment was divided by the sum of presynaptic terminals −1 in that segment, with the following equation:

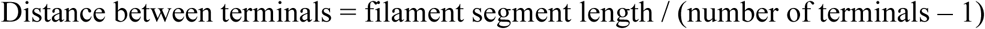

Terminal distance measurements for filaments comprising one neuron were then averaged to create a single data value for each neuron. Presynaptic terminal diameter values were exported directly from Imaris as an Excel file, where each terminal is one data point.

### Statistics

Values and statistical tests are reported in each section and/or figure legend. All numbers are represented as the mean±SD unless otherwise noted. For all statistical analyses, datasets were first assessed for normality using the Shapiro–Wilk test. For comparisons among multiple groups when the data did not meet the assumptions for a parametric test, a Kruskal–Wallis test was performed, followed by Dunn’s post hoc test with Bonferroni correction for pairwise comparisons. For comparisons between two independent groups that did not meet the assumptions for a parametric test, a Mann–Whitney test was used. A one-way ANOVA was performed for comparisons among multiple groups when parametric assumptions were met, followed by Tukey’s post hoc test for pairwise comparisons. Statistical analysis was performed in either R (R Core Team, 2017), Microsoft Excel (Redmond, WA), or Prism (GraphPad, La Jolla, CA). For all box plots (unless noted otherwise in figure legend), the box represents the interquartile range, line indicates the median, and whiskers indicate the range, excluding outliers (which are shown). Dots overlaid onto box plots generally represent the individual experimental datapoints and are described in each figure as needed. For Figures 2 and 3, each dot overlaid on the graph represents the average of several individual FRAP experiments completed at different synaptic terminals (up to 5) across one neuron’s axonal arbor. Since multiple terminals were measured within each given neuron, sample sizes are provided as the number of individual experiments done within the number of different neurons that were sampled across the number of different larvae.

## Data Availability

Data/sets needed to evaluate the conclusions in this paper are either provided in the paper, the Supplementary Materials, or will be archived at Zenodo (DOI: 10.5281/zenodo.23677439).

Requests for DNA constructs should be submitted to the corresponding author.

## Author contributions

WI designed and conducted the majority of experiments, with significant help from SMA and KN. KLS helped to develop and implement the longitudinal FRAP approach. NB, WI, SMA, and KN generated DNA constructs and developed analysis tools. KN and WI designed and conducted the synaptic terminal size and distribution analysis. TW and VU designed the experiments in collaboration with WI and provided mentoring on the project. WI and TW wrote the manuscript, with significant contributions from SMA, VU, and KN. All authors read, edited, and approved the final manuscript.

## Acknowledgements

We would like to thank: Joseph Lebowitz for generously sharing his inverted confocal microscope; Pamela McLean for α-synuclein DNA; Teresa Nicolson for p5E-neurod; Alex Nechiporuk and Kara Cerveny for sharing reagents and embryos; Leah Weston, Zac Tobias, and Greta Glover for reagent generation and protocol assistance; and Norma Velazquez-Ulloa and Luan Doster for helpful discussions.

