## Supplementary material for "The Parkinson’s disease-associated A53E mutation decreases alpha-synuclein aggregation and phosphorylation *in vivo*": NA

**Supplementary Information**

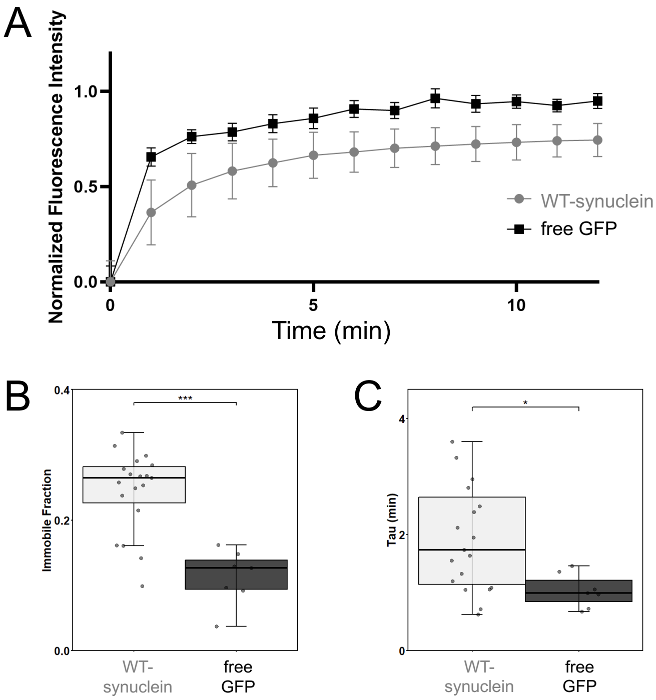


**Supplementary Figure 1.** Mobility of WT-α-synuclein differs from GFP at 4 days post fertilization (dpf). A) Normalized fluorescence recovery curves show the average fluorescence intensity over time following photobleaching for neurons expressing free GFP (black squares, n=12 experiments from 7 neurons from 6 larvae) or WT-α-synuclein (gray circles, n=50 experiments from 19 neurons from 16 larvae). B) The immobile fraction (IF) of WT-α-synuclein-GFP is significantly greater than free GFP (WT: 0.24±0.06; free GFP: 0.11±0.04; Mann-Whitney p=0.0002). C) Recovery tau for WT-α-synuclein is significantly greater than free GFP mobility (WT: 2.1±1.5 min; free GFP: 1.03±0.3 min; Mann-Whitney p=0.01). For B and C, boxes represent the interquartile range, line indicates median, and whiskers indicate range, with individual neuron measurements overlaid. For B-C, each dot overlaid on the graph represents the average of several individual FRAP experiments completed at different synaptic terminals within one neuron's axonal arbor. Statistical significance indicated by * p<0.05, ***p<0.005.

Data table from longitudinal FRAP experiments

| Condition | Neuron ID | 2 dpf IF | stdev | n (# experiments within this neuron at this age) | 4 dpf IF | stdev | n | 6 dpf IF | stdev | n |
|---|---|---|---|---|---|---|---|---|---|---|
| WT | 1 | 0 | N/A | 2 | 0.161316 | 0.08187 | 3 | 0.374697 | N/A | 2 |
| WT | 2 | 0.229152 | 0.01861 | 4 | 0.253009 | 0.01954 | 3 | 0.617005 | 0.1153 | 4 |
| WT | 3 | 0.191 | 0.1093 | 3 | 0.2842 | 0.148 | 3 | 0.4704 | 0.07554 | 3 |
| E46K | 1 | 0.275538 | 0.1115 | 3 | 0.597215 | 0.2254 | 3 | 0.445742 | 0.1664 | 3 |
| E46K | 2 | 0.295551 | N/A | 2 | 0.433038 | 0.09783 | 4 | 0.657302 | 0.2745 | 3 |
| E46K | 3 | 0.293511 | N/A | 2 | 0.406969 | 0.1628 | 4 | 0.535568 | 0.1109 | 4 |
| E46K | 4 | 0.246198 | 0.1995 | 4 | 0.387071 | 0.03494 | 3 | 0.461438 | 0.1617 | 3 |
| E46K | 5 | 0.3015 | 0.06955 | 3 | 0.367 | 0.1466 | 3 | 0.5193 | 0.1247 | 4 |
| A53E | 1 | 0.185741 | 0.1098 | 3 | 0.133278 | 0.1154 | 3 | 0.19981 | 0.04887 | 3 |
| A53E | 2 | 0.179486 | 0.139 | 4 | 0.269818 | 0.1213 | 3 | 0.284377 | 0.1372 | 4 |
| A53E | 3 | 0.1502 | 0.1347 | 3 | 0.229 | 0.1226 | 3 | 0.1667 | 0.1427 | 3 |
| A53E | 4 | 0.1722 | 0.1632 | 3 | 0.308 | 0.05287 | 5 | 0.221 | 0.03627 | 3 |
| A53E_S129D | 1 | 0.2114 | 0.1225 | 4 | 0.3764 | 0.07428 | 3 | 0.5934 | N/A | 2 |
| A53E_S129D | 2 | 0.04753 | N/A | 2 | 0.3238 | N/A | 2 | 0.2895 | 0.0538 | 3 |
| A53E_S129D | 3 | 0.2882 | N/A | 2 | 0.3139 | N/A | 2 | 0.5233 | N/A | 2 |
| A53E_S129D | 4 | 0.1958 | N/A | 2 | 0.34 | N/A | 2 | 0.4075 | 0.07792 | 3 |

**Supplementary Figure 2.** This table provides the average immobile fraction (IF) values from individual neurons that were measured longitudinally from 2 to 6 days post fertilization (dpf). Each average IF value represents multiple synaptic terminals imaged in the same individual neuron (see n column). Standard deviation is reported for n>2.

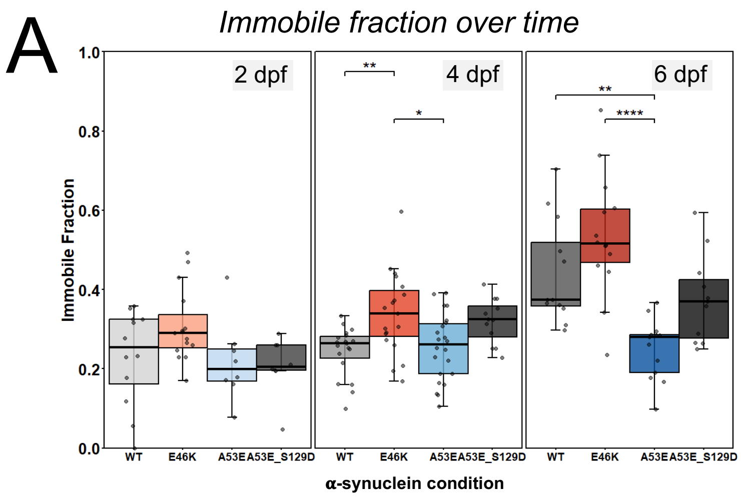


**Supplementary Figure 3.** Genetic modification to mimic phosphorylation in A53E-α-synuclein is sufficient to decrease its mobility over time *in vivo*. In order to compare A53E+S129D directly to the other conditions, some data are repeated from Figure 3 and 6B. A) *In vivo* FRAP experiments quantified immobile fraction measured at 2, 4, and 6 days post fertilization (dpf), grouped by age. Data is repeated from Figure 6B but rearranged for within-age comparison. Kruskal Wallis 4 dpf $p<0.003$; Dunn's multiple comparison 4 dpf WT vs. E46K $p=0.01$, 4 dpf E46K vs. A53E $p=0.04$. Kruskal Wallis 6 dpf $p<0.0001$; Dunn's multiple comparison 6 dpf WT vs. A53E $p=0.004$, 6 dpf E46K vs. A53E $p<0.0001$. Boxes in represent the interquartile range, line indicates median, and whiskers indicate range. Each dot overlaid on the graph represents the average of several individual FRAP experiments completed at different synaptic terminals within one neuron's axonal arbor. Statistical significance indicated by * $p<0.05$, ** $p<0.01$, and **** $p<0.0001$.
